# DNA2Graph enables automated identification of non-linear DNA molecules in electron microscopy

**DOI:** 10.64898/2026.08.05.742986

**Authors:** Federico Chinello, Michele Giannattasio, Elia Zanella, Flavia Bruno, Francesca M. Buffa, Ylli Doksani

**Affiliations:** IFOM ETS – The AIRC Institute of Molecular Oncology, Milan, 20139, Italy; Department of Computing Sciences and Bocconi Institute for Data Science and Analytics, Bocconi University, Milan, 20136, Italy; Dipartimento di Oncologia ed Emato-Oncologia, Università degli Studi di Milano, Via Festa del Perdono 7, 20122 Milan, Italy

## Abstract

Electron microscopy of spread and rotary-shadowed DNA molecules provides a direct readout of DNA structure and remains uniquely informative for studying replication, recombination and repair intermediates. However, quantitative EM analysis is limited by the rarity of biologically informative structures and by the time required for expert operators to inspect very large numbers of molecules. Automated image acquisition and stitching have increased the scale of EM datasets but have shifted the main bottleneck from image collection to image analysis. Standard image segmentation tools and machine-learning approaches fail to faithfully preserve molecular continuity in EM images of DNA molecules contrasted by rotary shadowing.

Here we present DNA2Graph, an open-source software for segmentation of DNA molecules from EM images. Rather than treating segmentation as a purely pixel-level task, DNA2Graph represents each molecule as a spatial graph of nodes and edges. It applies dedicated graph-based error-correction algorithms that repair signal interruptions and spurious connections introduced during segmentation, while enforcing the biological priors of continuity and thinness.

DNA2Graph classifies molecules as linear or non-linear, thereby converting large imaging datasets into focused lists of candidate structures for operator review. We validated DNA2Graph on two genomic DNA datasets: a structure-poor, non-enriched human sample and a structure-rich yeast sample. DNA2Graph reduced the number of molecules requiring operator review by 45-fold and 7-fold, respectively. In addition, DNA2Graph-assisted review slightly improved the operator’s structure-detection sensitivity compared to unaided image review.

The software also enables automated length measurement of individual DNA molecules and generates machine-readable outputs for downstream quantitative and computational analyses. DNA2Graph does not require training on manually annotated data, can run on a personal computer and can be adapted across experimental preparations through interpretable parameters.

## Introduction

DNA structural intermediates, such as Y-shaped replication forks and X-shaped Holliday junctions, are defining features of DNA replication, recombination and repair. Their direct visualization can provide unique mechanistic insights into the pathways that preserve genome integrity. The most direct way to monitor these intermediates is visualization by electron microscopy (EM). In order to reveal their molecular contour, DNA filaments are forced to assume a 2D configuration via spreading as monomolecular layers on an aqueous surface. The first EM spreading methods relied on the capture of DNA filaments by a protein monolayer, typically cytochrome c, at an air-water interface, whereas later variants replaced the protein layer with a cationic surfactant, benzyldimethylalkylammonium chloride (BAC) (1, 2). Spread DNA molecules are then transferred by contact onto carbon-coated grids pre-treated to promote DNA adsorption and are subjected to metal rotary shadowing to reveal the molecular contour (3, 4). These early single-molecule imaging techniques enabled direct visualization of DNA molecules engaged in replication or recombination (5-9). Over the years, EM imaging has also documented additional DNA structures, including extrachromosomal circular DNA in higher organisms, rolling-circle replication intermediates, telomeric t-loops that protect chromosome ends from the DNA damage response, i-loop structures, formed during intramolecular recombination at repetitive elements and R-loops formed during transcription (10-15). In addition to providing direct visualization, quantitative analysis of structural intermediates can reveal how genetic pathways or cellular conditions shape DNA metabolism. For example, EM analysis showed that checkpoint-defective cells accumulate reversed or processed replication forks, revealing the nature of their replication defects (16). Similarly, visualization of ssDNA flaps at replication forks identified these structures as possible substrates of the DNA2 nuclease (17).

A fundamental limitation of quantitative EM analysis is the paucity of structural intermediates relative to linear DNA molecules. Depending on the organism, cell-cycle state and experimental preparation, replication or recombination intermediates are hundreds-fold less abundant than linear DNA molecules, while individual subclasses of structures can be substantially rarer. This low frequency limits statistical power and restricts the applicability of EM analysis, especially for rare structural intermediates. To mitigate this limitation, EM analysis is often preceded by enrichment steps designed to increase the fraction of molecules carrying structures of interest. However, these procedures are technically challenging, require large amounts of starting material and may introduce biases in the types of structures recovered (18-20). Even in enriched samples, identification of structural intermediates remains time-consuming because an operator must visually inspect thousands of molecules per sample. More recently, automated image acquisition and stitching have substantially increased the size of EM datasets by enabling the collection of large, contiguous imaging areas (21). As a result, the major bottleneck has shifted from image collection to image analysis. Here we report the development of DNA2Graph, an open-source image-analysis software for EM images of DNA molecules visualized by rotary shadowing. DNA2Graph implements a computer-vision workflow to: i) segment DNA molecules in large, stitched EM images; ii), represent them as spatial graphs, classify them as linear or non-linear candidate molecules, and iii) provide automated length measurements of individual molecules. By converting large EM datasets into focused lists of non-linear candidate molecules, DNA2Graph reduces the number of molecules requiring visual inspection. In the validation datasets, this reduction was 45-fold for a structure-poor non-enriched human sample and 7-fold for a structure-rich yeast sample. This reduction in the number of molecules that require visual inspection is estimated to save from 1.2 to 5.7 hours per sample, in the task of recovering a standard-sized set of DNA structures. To our knowledge, DNA2Graph is the first tool that provides: i) automated, continuity-preserving segmentation of individual DNA molecules visualized by rotary shadowing; ii) their classification into linear and non-linear candidates; iii) per-molecule length measurement. DNA2Graph therefore addresses a major practical bottleneck in EM-based analysis of DNA structural intermediates, supporting both laboratories that routinely perform single-molecule analysis of DNA by EM and research groups that access this technique through collaborations with specialized electron microscopy units. In addition, its graph-based output provides a machine-readable representation of DNA molecules for downstream quantitative and computational analyses.

## Materials and methods

We release DNA2Graph as an open-source Python package and make its source code publicly available at https://github.com/chinefed/DNA2Graph. Comprehensive documentation is also provided, including a technical manual describing the internal algorithms of DNA2Graph (available as Supplementary Data) and the DNA2Graph website (https://federicochinello.com/DNA2Graph/), which contains additional resources such as an installation guide and a series of video tutorials.

### Supported image formats

DNA2Graph supports 8-bit and 16-bit grayscale images in TIFF format. DNA2Graph has been extensively tested on images acquired at pixel sizes of approximately 1.0-1.7 nm/pixel. This range is particularly well suited to the acquisition of large, stitched images, representing a good compromise between image resolution, acquisition time, and image size while still allowing a large grid area to be covered. Large deviations from this range may require parameter adjustment for optimal performance.

### Conventional image segmentation (Figure 1)

To illustrate the limitations of pixel-level segmentation on rotary-shadowed DNA (Figure 1), a conventional segmentation pipeline was applied in Fiji (ImageJ v1.54p) (22), independently of DNA2Graph. EM images were converted to 8-bit with contrast enhancement (0.35% saturated pixels) applied prior to conversion. Images were denoised with a median filter (radius = 2 px). DNA filaments were segmented by manual thresholding. An experienced user iteratively adjusted the grayscale intensity threshold and selected a value of 139 on the 8-bit scale as the threshold that best delineated filaments from the background. The resulting thresholded regions were converted to a binary mask. The mask was subjected to one round of morphological dilation followed by a closing operation to bridge small gaps. Segmented objects were then filtered using Analyze Particles, retaining objects ≥250 px^2^ with a circularity of 0.00-0.20 to select elongated filaments while excluding compact contaminating particles. For visualization, each retained ROI was assigned a distinct color, overlaid on the original micrograph, and the composite image was flattened to an RGB image.

**Figure 1.**
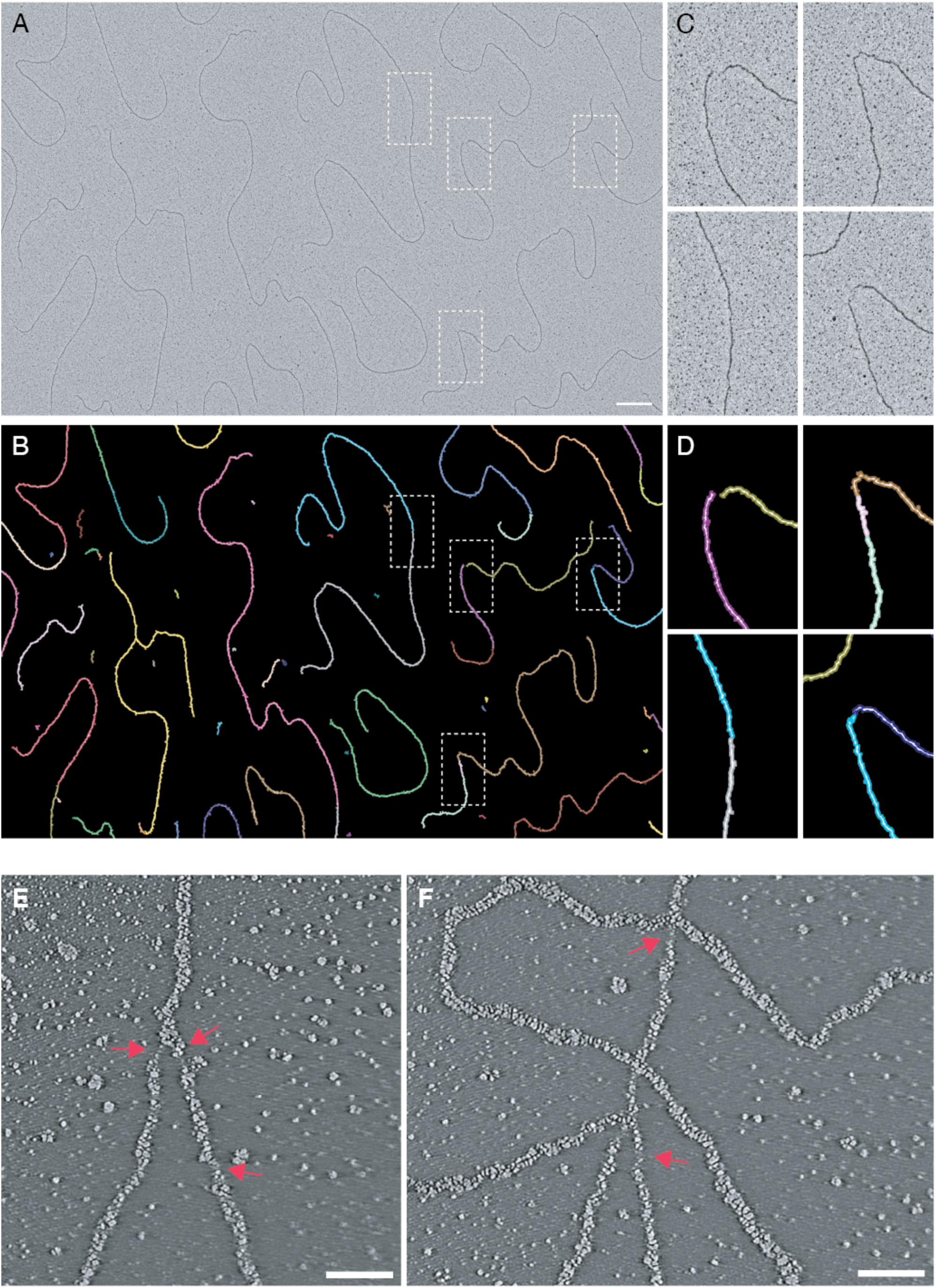
Intensity-based segmentation fails to recover continuous DNA molecules from rotary-shadowed electron micrographs. **(A)** Transmission electron micrograph of rotary-shadowed DNA, with an optimal spreading and rotary shadowing quality and contrast. Individual molecules appear as dark, continuous filaments. Dashed boxes mark the regions magnified in (C). Scale bar, 500 nm. **(B)** Output of classical threshold-based segmentation of the micrograph in (A), using expert-optimized manual thresholding (see Methods). Each color denotes a distinct segmented object. An intact molecule appears as a single continuous color. **(C)** Magnified views of the boxed regions in (A) indicating points were a DNA molecule is split into separate objects by the segmentation. **(D)** Magnified views of the boxed regions in (B), corresponding to the identical regions shown in (A). Filaments that are visibly continuous in the raw data (C) are split by the segmentation into separate objects, seen here as colour changes and gaps. Because the input is of optimal quality and the threshold was set to its optimum by an experienced user, these interruptions represent an intrinsic limitation of intensity-based thresholding for the thin, low-contrast filaments produced by rotary shadowing. **(E, F)** High-resolution electron tomography of rotary-shadowed DNA molecules, shown as single reconstructed slices from two independent acquisitions. The DNA appears as a granular filament. Arrowheads indicate regions of locally sparse platinum deposition along the DNA filament. These regions produce gaps in the signal along the molecule even though they do not correspond to genuine discontinuities in the underlying DNA. Scale bars, 20 nm.

### Preparation of the 6.211 kb linear DNA fragment

A 6.211 kb plasmid was linearized with ScaI and the resulting linear fragment was purified from an agarose gel. The purified DNA was resuspended in 100 µL of TE 1X at 10 ng/µL and 4, 5′, 8-trimethylpsoralen (TMP) (trioxsalen, Sigma, cat. no. T6137, resuspended 2 mg/mL in DMSO) was added to a final concentration of 30 µg/mL. The DNA solution was deposited on a parafilm strip and irradiated for 1 minute on a Stratalinker 1800 (Stratagene, cat. No. 400072) carrying 365 nm UV bulbs at a distance of ∼5 cm from the bulbs. The DNA was then precipitated with sodium acetate and isopropanol and resuspended in 50 µL of TE 1X. 50 ng of psoralen-crosslinked DNA was spread by the BAC-formamide method (see DNA spreading and rotary shadowing).

### Length distribution analysis of the 6.211 kb control fragment (Figure 4)

Molecule length measurements in pixel were recovered from the DNA2Graph output CSV file and converted to nm, by multiplication with the pixel size of the images (1.0655 nm/pixel). Values were pooled across all stitched images of the dataset and only molecules classified as linear were retained for length analysis. Frequency distributions were generated in GraphPad Prism using a bin width of 25 nm. To exclude the short molecules, resulting mostly from missegmentation of dirt on the grid, a length cutoff of 1000 nm was applied. The main peak of the resulting distribution was then fitted with a single Gaussian by nonlinear regression (GraphPad Prism), yielding the mean and standard deviation reported in the text. The length-per-kilobase was calculated by dividing the Gaussian mean by the known fragment size (6.211 kb). The apparent stretching factor was expressed relative to the canonical contour length of B-form double-stranded DNA (0.34 nm/bp, i.e. 340 nm/kb).

### Yeast genomic DNA sample preparation

*Saccharomyces cerevisiae* cells were arrested in G1 with α-factor and harvested at 105 minutes after release into S phase at 25°C in YPD containing 0.033% v/v methyl methane sulphonate, with samples harvested at 105 minutes from the G1 release. Genomic DNA was subjected to in vivo psoralen crosslinking, extraction and enrichment of replication intermediates essentially as described (19). Briefly, harvested cells were washed and resuspended in cold water and incubated with TMP (10 µg/mL final) followed by irradiation with 365 nm UV light; the TMP addition and irradiation cycle was repeated four times to ensure efficient psoralen-mediated DNA interstrand crosslinking. Genomic DNA was then extracted by the CTAB method and digested with PvuI. Replication intermediates were enriched on a benzoylated-naphthoylated-DEAE-cellulose (BND-cellulose) column, exploiting the affinity of the resin for the single-stranded DNA regions more abundant in DNA replication and repair intermediates. Linear duplex molecules were preferentially eluted in 1 M NaCl and the RI-enriched fraction was eluted in 1M NaCl buffer containing 1.8% w/v caffeine. The eluate was purified, concentrated and subjected to a buffer change on a size-exclusion filter column before spreading.

### Human genomic DNA sample preparation

RPE1-hTERT cells were maintained under standard culture conditions. Cells were harvested and subjected to in vivo psoralen crosslinking and genomic DNA extraction as described (23). Briefly the cell suspension was treated with TMP (30 µg/mL) and irradiated with 365 nm UV light for four successive cycles. Cells were lysed in TNES buffer containing RNase A, treated with proteinase K, and the DNA was extracted with phenol-chloroform-isoamyl alcohol followed by one extraction with chloroform and precipitated with isopropanol, with gentle handling throughout to preserve high-molecular-weight DNA. The genomic DNA was digested with KpnI, which yields fragments of approximately 10 kb, and used directly as a non-enriched, structure-poor preparation for spreading.

### DNA spreading and rotary shadowing

DNA was spread and rotary-shadowed essentially as described (19, 24). Briefly, ∼30-50 ng of DNA was spread by the BAC method. The DNA was mixed with formamide and BAC and applied to a water hypophase using a freshly cleaved mica sheet as a ramp, forming a monomolecular DNA-detergent film at the air-water interface. The monomolecular DNA film was adsorbed onto carbon-coated copper grids produced by using electron beam evaporation technique. Before use, the carbon-coated grids were activated by contact with an ethidium bromide solution (33.3 µg/mL). Grids with adsorbed DNA were stained with uranyl acetate dissolved at 0.2 µg/µL in absolute ethanol and subjected to rotary shadowing with 8 nm of platinum using the MED020 evaporator (electron beam evaporation) fitted with the low-angle rotary-shadowing kit, with the sample held at a shallow angle (∼3°) to the platinum source.

### Manual annotation by expert operators

Manual identification of DNA structures was performed by two expert operators using the ImageJ macro described previously (24). For each candidate, the operator drew a rectangular ROI containing the molecule identified as a structure and the macro automatically saved the set of ROIs for each stitched image. A structure was defined as a physical overlap or contact of separate DNA molecules or of distinct regions within the same molecule, or a circular DNA molecule.

The two operators worked independently and were blind both to each other’s annotations and to the DNA2Graph classification. Each operator first inspected the images and selected all objects meeting the structure definition as ROIs. Only after these ROIs had been generated was the operator given the second part of the task (i.e. comparison with the DNA2Graph output, as described below) so that the initial manual annotation was performed without knowledge of the software’s calls.

For each sample, all stitched images were analyzed in a single, uninterrupted session. A timer was started at the beginning of the session, and the per-stitch processing time was obtained by dividing the total annotation time by the number of stitched images.

### Performance metrics and time-saving estimates

To compare manual and DNA2Graph-assisted analysis, the operator ROIs and the DNA2Graph non-linear ROIs (candidate structures) were loaded onto the same stitched image. Each ROI was then classified into one of the following categories: found by DNA2Graph (all DNA2Graph calls); found by the operator (all operator annotations); overlapping (structures identified by both); operator only (structures identified by the operator but not by DNA2Graph); DNA2Graph only (non-linear ROIs reported by DNA2Graph but not identified by the operator); and false positives (DNA2Graph non-linear ROIs that did not meet the definition of a structure). A DNA2Graph-only object was scored as a real structure when it met the structure definition and as a false positive otherwise.

Ground truth was defined as the union of real structures identified by either approach (operator alone or DNA2Graph-assisted review). This definition was adopted because the DNA2Graph-assisted review identified additional structures that were not detected during the unaided operator review. Sensitivity (recall) was calculated as the fraction of ground-truth structures recovered by each method. The false discovery rate of DNA2Graph was expressed relative to the total number of non-linear objects it called (real structures plus false positives). False negative rates were expressed relative to the ground-truth set.

For the analysis of the full datasets (100 stitched images per sample), the DNA2Graph non-linear ROIs were loaded and visually inspected by the operator, and each was classified according to its structure type. Time-saving estimates were derived by comparing the time required to manually inspect all molecules across the full dataset with the time required to inspect only the DNA2Graph-selected non-linear candidates, using each operator’s measured per-stitch processing time (see Fig. 6 legend for the full calculation).

### EM image acquisition

EM images were acquired on a Talos F200C G2 transmission electron microscope (Thermo Fisher Scientific) operated in bright-field mode at 200 kV and equipped with a Ceta-S camera (an indirect-detection CMOS sensor of 4096 × 4096 pixels, physical pixel size 14 µm × 14 µm). For each DNA sample, one or more 30 µm × 30 µm surfaces were imaged as non-binned tiles with 10% overlap using MAPS software (Thermo Fisher Scientific), the number depending on the density of molecules on the grid and the type and number of structures to be analyzed. The pixel sizes most frequently used – chosen as a compromise between acquisition time, resolution and dataset size – were 1.0, 1.3, and 1.7 nm. The tiles covering each 30 µm × 30 µm surfaces were stitched in MAPS and exported as 16-bit TIFF files for downstream analysis with DNA2Graph.

As an example, at a pixel size of 1.3 nm, approximately 64 tiles (10% overlap) were required to cover a single 30 µm × 30 µm surfaces, generating a stitched TIFF of roughly 1.7 GB; acquiring 100 such surfaces (approximately 6, 400 tiles) took roughly 12–14 hours.

### EM tomography acquisition for Figure 1

A tilt series of a rotary-shadowed DNA sample was acquired on the Talos F200C G2 transmission electron microscope (Thermo Fisher Scientific) operated in bright-field mode at 200 kV and equipped with the Ceta-S camera, using the Thermo Fisher Tomography software 5 Images were recorded over a tilt range of ±54° at 1° increments following a dose-symmetric scheme, at a pixel size of 0.5 nm and a defocus of -1.86 µm. Tilt-series alignment was performed using fiducial-less patch tracking method of the Inspect3D software Thermo Fisher Scientific. Tomogram reconstruction was done with Inspect3D Thermo Fisher Scientific software using the SIRT algorithm (20 iterations) yielding a volume of 1472 × 1396 × 177 voxels (∼177 voxels thick). The reconstruction was visualized with Fiji (ImageJ).

## Results

### Faithful preservation of molecular continuity is the central challenge in segmenting rotary-shadowed DNA

A critical requirement for segmenting DNA molecules in EM images is the faithful preservation of molecular continuity. Because DNA filaments are extremely thin and long, even a small number of misclassified pixels can fragment a single molecule into multiple disconnected components (Figure 1A-D) or spuriously connect distinct molecules into an apparent structure. This problem is further complicated by the contrasting method. Rotary shadowing does not generate a perfect, continuous outline of the DNA filament; it produces a granular metallic replica (Figure 1E, F). The same granularity has two practical consequences for image analysis. In regions away from DNA, metal particles form a dense particulate background whose size and contrast can partially overlap with the DNA signal. Along the DNA itself, local variation in metal deposition can generate apparent signal interruptions even when the underlying molecule is physically continuous (Figure 1E, F).

A segmentation-only workflow is therefore trapped between two related connectivity errors. When the initial segmentation is too conservative, faint portions of a molecule are lost and a single DNA molecule is represented as several disconnected fragments. Relaxing the segmentation can recover some weakly contrasted regions, but it cannot recover stretches where the rotary-shadowed replica contains no signal despite the underlying DNA molecule being continuous. At the same time, a more permissive segmentation increases the incorporation of background metal particles and close molecular contacts, causing distinct molecules to be merged into false branched or cyclic objects. This limitation is not restricted to simple thresholding: CNN-based segmentation approaches can also suffer from local discontinuities and molecule fragmentation, especially in poorly contrasted regions (21).

Standard morphological operations, which clean binary masks by expanding, shrinking or closing gaps between segmented pixels, cannot fully solve the problem, because gaps that interrupt a molecule can be comparable in size to gaps that separate distinct molecules. Thus, a mask that appears locally accurate can still be unusable for downstream analysis if it breaks one molecule into fragments or merges neighboring molecules into false structures. For rotary-shadowed DNA, segmentation quality must therefore be judged not only by pixel overlap with an annotation, but also by the preservation of the correct molecular continuity.

### DNA2Graph segments DNA while preserving molecular continuity

We developed DNA2Graph to address this molecular continuity problem directly. The software accepts one or more grayscale EM images as input and can be run either through a graphical user interface for routine use on a personal computer, or through a command-line interface for scalable processing on high-performance computing clusters.

The DNA2Graph workflow has two conceptual stages (Figure 2 and Supplementary data). First, a segmentation module generates a raw binary mask using an image-processing pipeline we engineered to be robust across diverse inputs. This pipeline suppresses background noise, enhances the DNA signal, and preserves continuity in segmented DNA molecules while minimizing false merging between DNA molecules and noise-induced artifacts. Second, this raw mask is converted into a spatial graph, in which nodes represent points along each segmented molecule and edges encode connectivity. Error correction is then performed on the graph rather than on the image alone. This graph-based representation allows the software to ask whether two signal fragments are likely to belong to the same continuous molecule, whether a contact should be resolved, and whether an object is compatible with the expected thin, filamentous geometry of DNA. The key correction step restores molecular connectivity by selectively adding graph edges between candidate nodes that are close in image space but distant, or disconnected, in graph space. This allows DNA2Graph to bridge local signal interruptions compatible with a continuous molecule, rather than simply closing all nearby gaps in the mask. The resulting graph is then refined to remove residual artifacts (detailed in Supplementary data). Once the corrected graph is generated, DNA2Graph renders it back into a refined binary mask and exports the corresponding molecular objects for visualization and downstream measurement (Figure 2). In this way, segmentation is used as an entry point, but the final molecular representation is determined by graph-based correction rather than by pixel classification alone (Figure 3).

**Figure 2.**
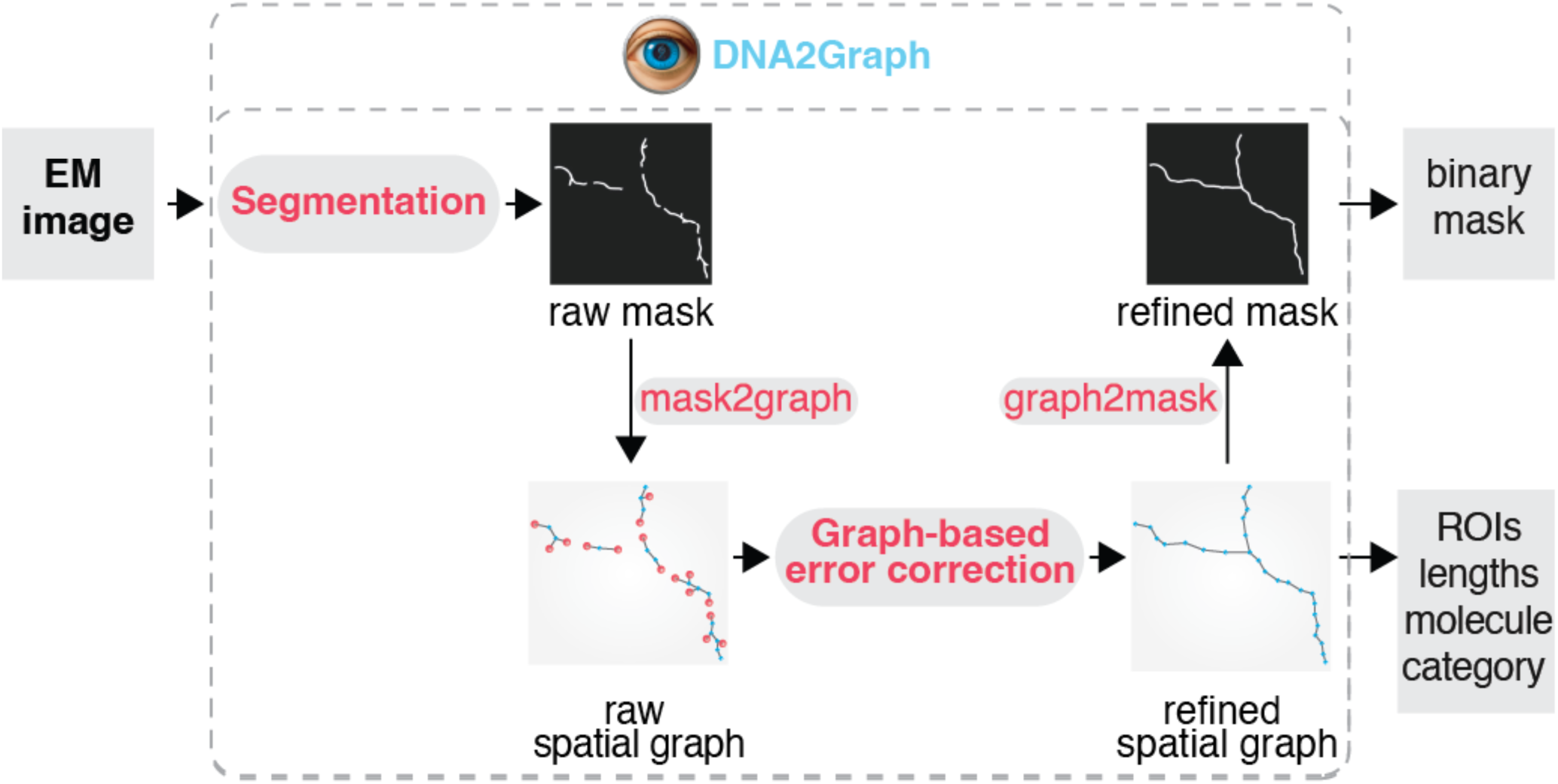
The DNA2Graph workflow. Schematic overview of the DNA2Graph pipeline. An input EM image is segmented into a binary mask (raw mask), which is converted into a spatial graph representation (mask2graph). Graph-based error correction reconnects fragmented segments, whose endpoints are shown as red nodes in the raw spatial graph, yielding the refined spatial graph. The corrected graph is converted back into a binary image (graph2mask) to produce the refined mask. DNA2Graph outputs the refined binary mask and, from the corrected graph, ImageJ ROIs, molecule length and molecule categories (“boundary”, “linear”, “non-linear”).

**Figure 3.**
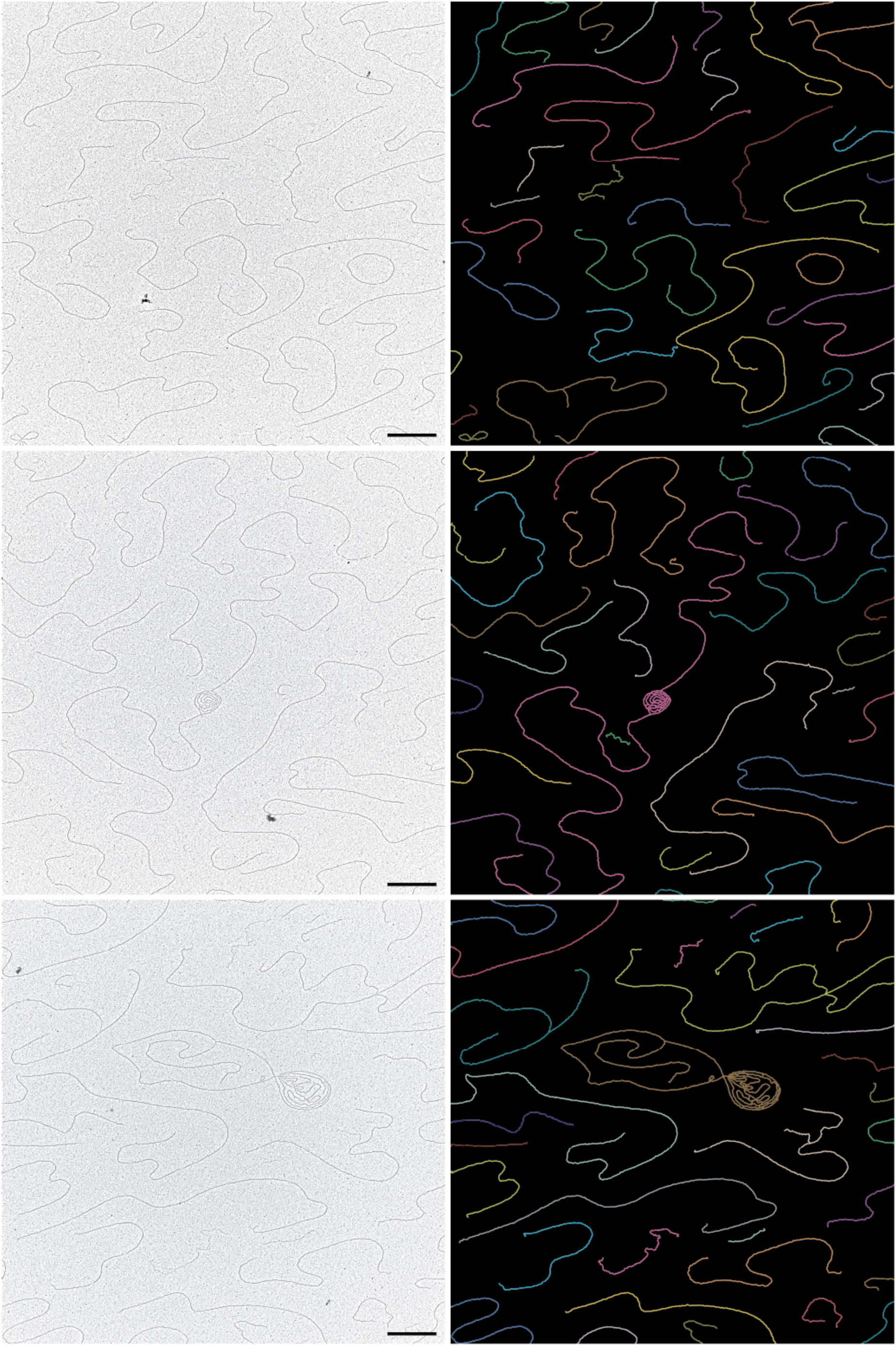
DNA2Graph recovers continuous single-molecule objects. Three representative rotary-shadowed EM micrographs (left) and their corresponding DNA2Graph mask output (right). The ROIs of the segmented molecules in the mask output are shown as colored lines. Each color denotes a distinct ROI. Because graph-based error correction reconnects fragmented segments, individual DNA molecules are recovered as single continuous ROIs, each represented by one color along its full length. This is in direct contrast to the intensity-based segmentation method shown in Figure 1, where a single molecule was split into several differently colored fragments. Scale bars, 1000 nm.

To test whether DNA2Graph effectively preserves molecular continuity and supports length measurements, we analyzed a dataset of stitched EM images containing a homogeneous population of a 6.211 kb linear restriction fragment (Figure 4A). DNA2Graph identified 7, 114 linear molecules, whose lengths showed a major peak near 2600 nm, with 75% of molecules falling between 2400 and 2800 nm, together with a minor population below 1000 nm (Figure 4B). Visual inspection of the top 100 longest molecules, ranging from 3268-5137 nm, confirmed that they were all correctly segmented long DNA molecules, likely bacterial genomic DNA contamination. Visual inspection of 501 of the 1, 103 molecules shorter than 1000 nm showed that they did not arise from discontinuities in the segmentation of linear molecules, but mostly corresponded to contaminating particles, local grid artifacts and some correctly segmented shorter DNA molecules (Figure 4C-E). Applying a conservative 1000 nm length cutoff removed most of these residual objects, after which 89% of molecules fell within the 2400-2800 nm range (Figure 4F). Gaussian fitting of the main peak gave a mean ± SD of 2631 ± 65 nm, corresponding to 423.6 ± 10.5 nm/kb for the 6.211 kb fragment (Figure 4F). Compared with the canonical contour length of B-form dsDNA, this corresponds to an apparent stretching factor of 1.246 ± 0.030 (Figure 4G). This value is likely to be a more accurate estimation of DNA stretching in BAC-formamide spreads, because DNA2Graph enables a more accurate tracing of individual DNA molecules, compared to manual tracking that typically shows an apparent stretching factor of 1.059 (360 nm/kb). Together, these results show that DNA2Graph preserves the continuity of linear DNA molecules while maintaining a low rate of inappropriate molecule joining.

**Figure 4.**
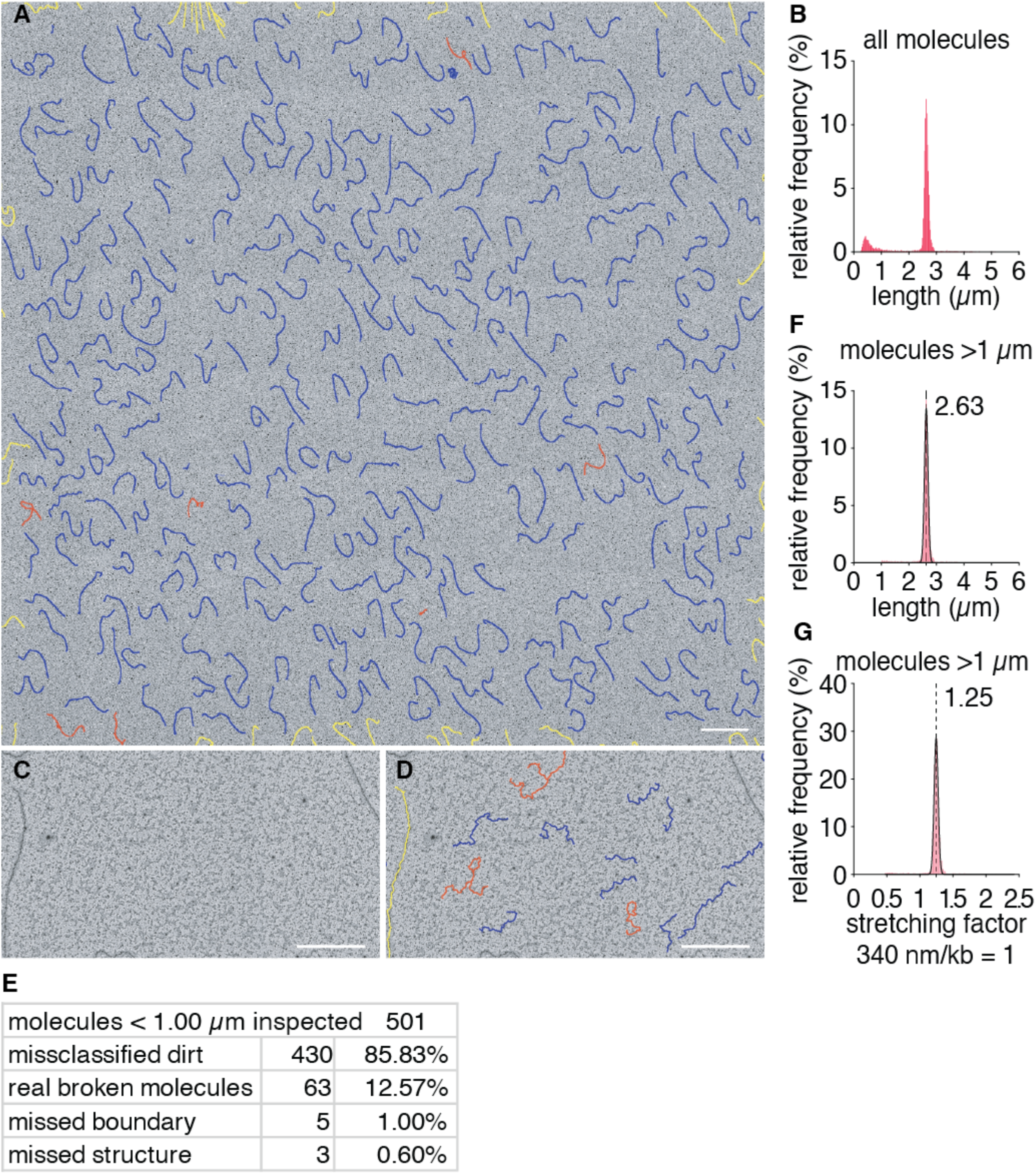
DNA2Graph preserves molecular continuity and enables accurate length measurement of a homogeneous DNA population. **(A)** Representative field of view from a stitched EM dataset of a homogeneous sample of a 6.211 kb restriction fragment, segmented by DNA2Graph. Each segmented molecule is drawn as a colored line according to its classification: blue, linear molecules; red, structures; yellow, molecules touching the field boundary. Scale bar, 2000 nm. **(B)** Length distribution of linear molecules segmented from 20 large fields as in (A) (n = 7, 114). The distribution shows a major peak around 2600 nm and a minor population near 500 nm. **(C)** Example of a region on the grid showing particulate dirt deposition. Scale bar, 500 nm. **(D)** Segmentation of the region in (C), showing that particulate contamination is segmented as short spurious objects. Scale bar, 500 nm. **(E)** Results of visual inspection of a subpopulation of linear molecules shorter than 1000 nm (501 of 1, 103 inspected), classifying their origin. **(F)** Length distribution of linear molecules after applying a 1000 nm cutoff, from the same dataset as (B) (n = 6, 011). Solid curve, Gaussian fit (mean ± SD = 2631 ± 65 nm; R² = 0.99); dashed line, fitted mean. **(G)** Distribution of the apparent stretching factor for the molecules in (F), calculated as measured length divided by the expected B-form contour length of the 6.211 kb fragment. Solid curve, Gaussian fit (mean ± SD = 1.246 ± 0.030; R² = 0.99); dashed line, fitted mean.

### DNA2Graph reduces expert inspection by enriching for candidate structures

After graph correction, each molecular object is assigned to one of three operational classes. Linear molecules are simple non-branching filaments, whereas non-linear molecules include branched or cyclic objects such as replication forks, Holliday junction-like structures (in general four branched X-shaped DNA molecules), bubbles, loops or circular molecules. These non-linear objects are reported as candidate structures for expert inspection, without assigning them to a specific biological subclass. Boundary molecules, which contact the image border and are likely incomplete, are flagged separately and can be excluded from downstream analysis.

The classified objects are exported as ImageJ-compatible ROI files, allowing DNA2Graph calls to be overlaid on the original image for inspection or manual refinement (Figure 5).

**Figure 5.**
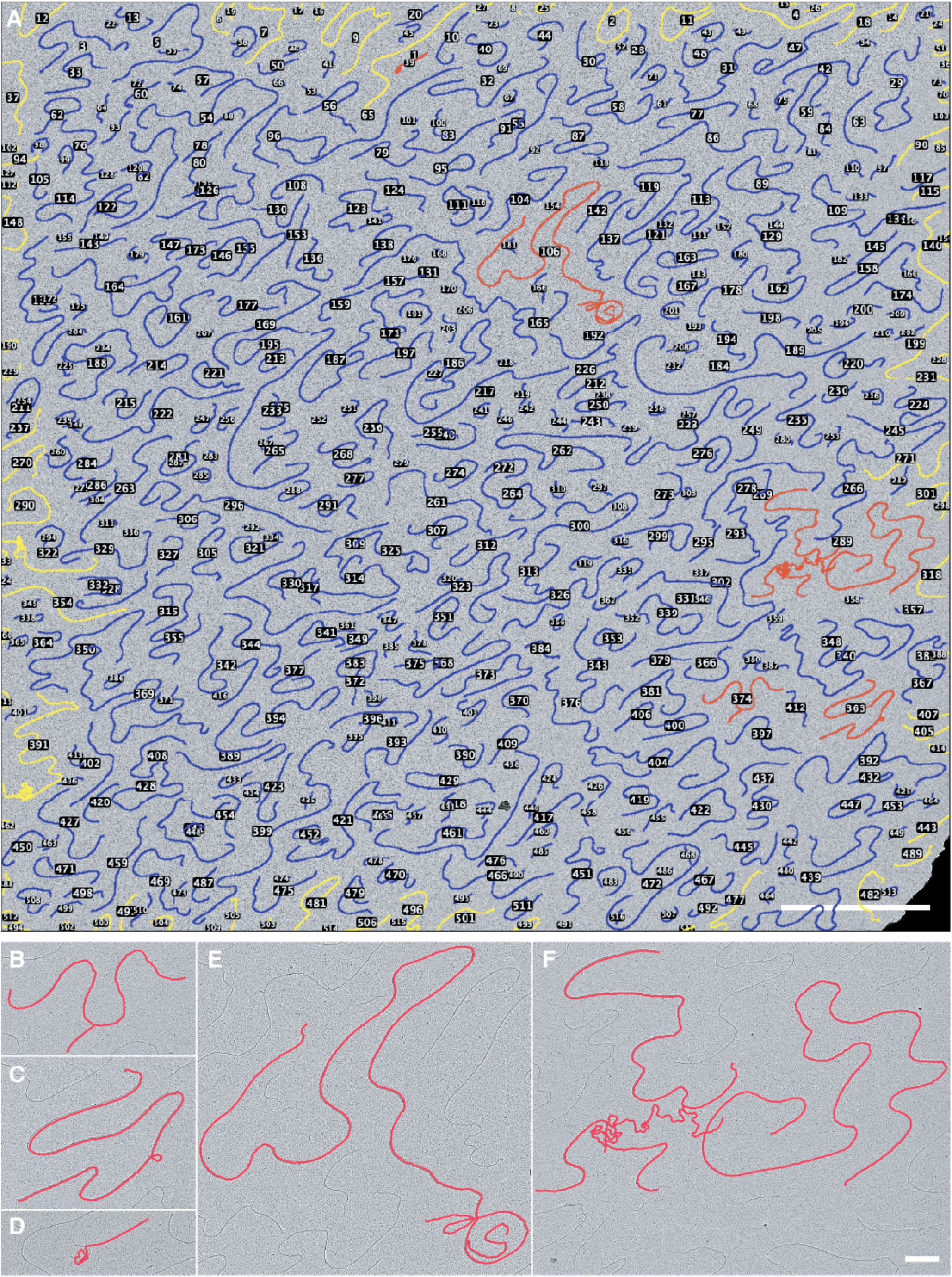
DNA2Graph enriches rare non-linear structures from a large EM field. **(A)** DNA2Graph segmentation of a large, stitched EM field. Each object is drawn as a colored trace according to its classification: blue, linear molecules; yellow, molecules touching the field boundary; red, non-linear structures. Of 517 total molecules identified, 434 were classified as linear, 78 as boundary, and 5 as non-linear. Numbers indicate individual object identifiers. Scale bar, 5000 nm. **(B-F)** Magnified views of the five non-linear molecules identified in (A). (B) A Y-shaped structure consistent with a replication fork. (C, E) Molecules containing internal loops. (D, F) Molecules with strand crossings arising from locally poorly spread (wiggly) filaments. The non-linear class comprises a candidate pool for expert triage rather than a set of confirmed structures: it contains both genuine biological candidates (B, C, E) and spreading artifacts (D, F), which the operator distinguishes on inspection. Scale bars, 500 nm.

The practical goal of DNA2Graph is not to replace expert interpretation of DNA structures, but to replace the exhaustive visual search required to find rare candidate structures. In a typical EM field, the large majority of molecules are linear and biologically uninformative for structure-focused analyses. In the segmentation of the large stitch (31.8 µm x 31.1 µm) shown in Figure 5A, DNA2Graph identified 517 molecules, 434 linear, 78 boundary and 5 non-linear molecules, which is the subclass that would be ultimately inspected by the operator for biologically relevant DNA structures. By automatically separating linear molecules from non-linear candidates and boundary objects, DNA2Graph converts the task from inspecting all molecules in a stitched image to inspecting a substantially enriched candidate list (Figure 5A-F).

To estimate the practical utility of DNA2Graph, we compared standard visual identification of DNA structures with DNA2Graph-assisted candidate selection in two genomic DNA datasets representing different levels of structure abundance. The first dataset consisted of non-enriched genomic DNA from unsynchronized human cells, a structure-poor condition expected to contain few replication or recombination intermediates. The second consisted of yeast genomic DNA from S-phase-synchronized cells, passed through a BND-cellulose column (18), a structure-rich condition containing a high frequency of replication intermediates. The full datasets consisting in 100 large, stitched images of 32 µm x 31 µm were processed with DNA2Graph, and representative subsets of 20 and 10 stitched images, respectively, were manually analyzed by two operators to compare the sensitivity of DNA2Graph in identifying DNA structures. Here, a structure is defined as a physical overlap or contact between separate DNA molecules on the grid, a contact between distinct regions of the same molecule, or a circular DNA molecule. This definition includes bubbles, Y-shaped molecules, X-shaped molecules, double Y termination structures, i-loop-like and t-loop-like molecules, and complex branched molecules. The DNA2Graph output was visually inspected by each operator and classified as a real structure (matching the definition above) or a false positive (arising from missegmentation of linear molecules). Using the union of real structures identified by either approach (operator alone or DNA2Graph-assisted operator) as ground truth, DNA2Graph-assisted review recovered a higher fraction of real structures than unaided expert analysis in both datasets and against both operators, exceeding each operator’s sensitivity by 1.5 to 4.8 percentage points (Figure 6A, B). Thus, an automated, deterministic analysis matched and modestly surpassed expert operators in detecting genuine DNA structures across two independent samples. The false discovery rate of DNA2Graph was sample-dependent: ∼50% in the structure-poor human dataset (sample 1) and ∼13% in the structure-rich yeast dataset (sample 2) (Figure 6A), largely due to dirt particles in proximity to linear molecules.

**Figure 6.**
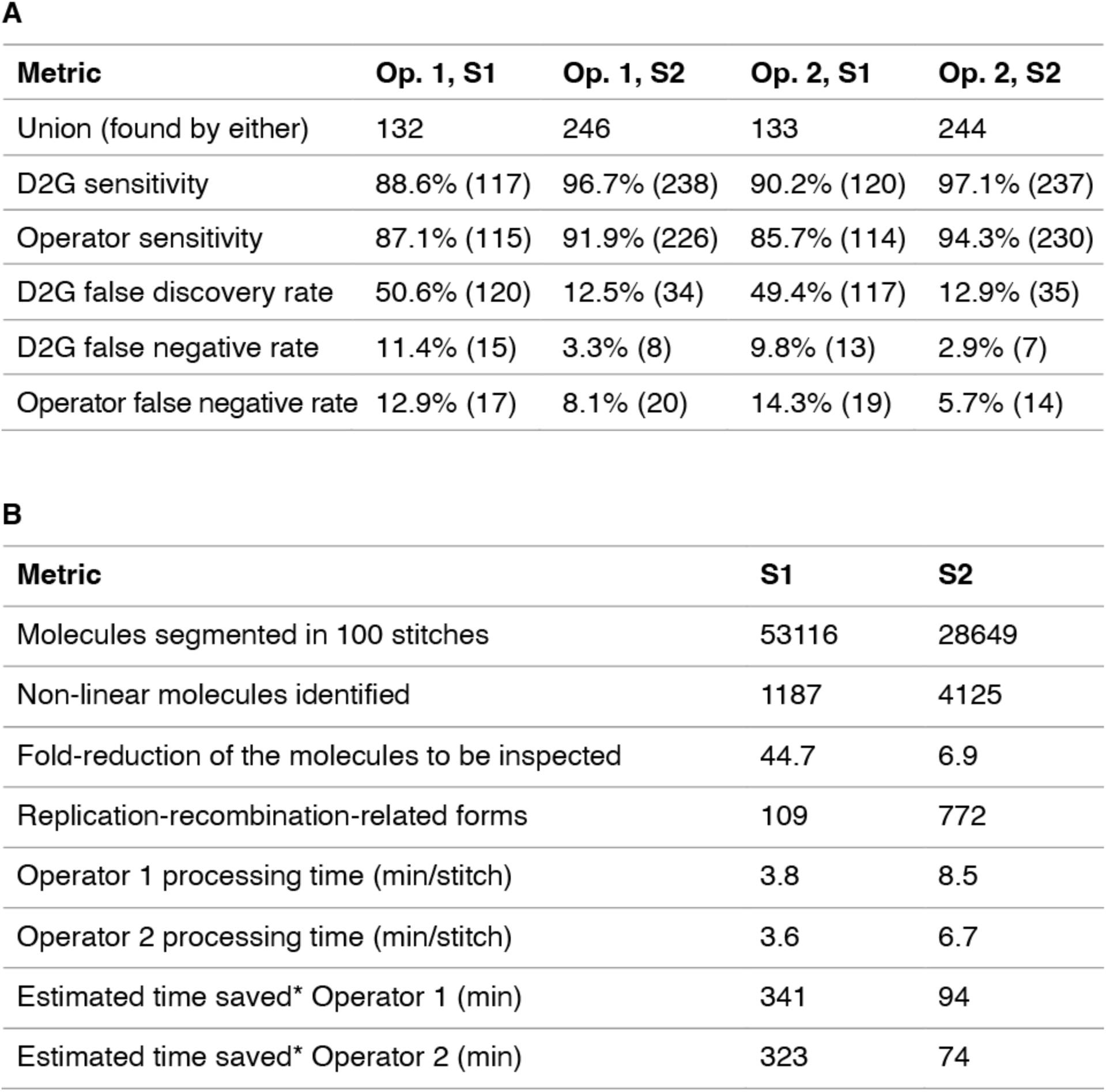
DNA2Graph matches operator sensitivity while substantially reducing inspection workload. Comparison of manual structure identification with DNA2Graph-assisted analysis in two datasets of differing structure abundance: sample 1 (S1; non-enriched genomic DNA from unsynchronized human cells, structure-poor) and sample 2 (S2; S-phase-synchronized yeast genomic DNA enriched for replication intermediates on a BND-cellulose column, structure-rich). **(A)** Performance of DNA2Graph relative to two independent operators (Operator 1 and Operator 2), each evaluated on a manually analyzed subset (20 stitched images for sample 1, 10 stitched images for sample 2). Ground truth was defined as the union of real structures identified by either the unaided operator or DNA2Graph-assisted inspection. Sensitivity (recall) is the fraction of ground-truth structures recovered. The false discovery rate is the fraction of DNA2Graph calls judged by the operator not to be real structures, relative to the total number of non-linear objects called by DNA2Graph in the subset (i.e. real structures plus false positives). The false negative rate is the fraction of ground-truth structures missed by each method, relative to the union. DNA2Graph sensitivity exceeded that of each operator in all four comparisons, by 1.5 to 4.8 percentage points. **(B)** Workload and time-saving estimates from running DNA2Graph on the full datasets (100 stitched images per sample). The fold-reduction in molecules requiring inspection is the ratio between all molecules identified by DNA2Graph and the non-linear molecules presented to the operator for inspection. *Estimated time saved was calculated as follows: the time to manually screen all molecules across the 100 stitches (100 × per-stitch processing time) was compared with the time to inspect only the DNA2Graph-selected non-linear candidates, obtained by converting the non-linear molecule count into stitch-equivalents (non-linear molecules identified / mean molecules per stitch) and multiplying by the same per-stitch time. The difference was normalized to the number of stitches required to accumulate 100 replication/recombination intermediates (assuming a typical analysis requires ∼100 such intermediates for sufficient statistical power). Values are given separately for each operator, using that operator’s measured per-stitch processing time.

When DNA2Graph was run on both full datasets (100 stitched images) the number of molecules requiring operator inspection (all molecules vs. the non-linear subset) was reduced ∼45-fold for sample 1 and ∼7-fold for sample 2 (Figure 6B). Visual inspection of the non-linear class (DNA2Graph-assisted analysis) yielded 109 molecules with shapes compatible with replication or recombination intermediates (Y-shaped, bubble-shaped, double-Y/termination, and X-shaped) in sample 1, and 772 such molecules in sample 2. Assuming a typical analysis requires ∼100 of these structural intermediates for sufficient statistical power and given each operator’s measured screening time per stitched image, DNA2Graph-assisted analysis is estimated to save, on average, ∼330 minutes of focused manual inspection in the structure-poor human dataset and ∼85 minutes in the structure-rich yeast dataset (Figure 6B).

These estimates are likely conservative, as they do not account for reduced operator performance during prolonged visual inspection and they scale linearly with the number of samples analyzed. Thus, DNA2Graph does not replace expert interpretation, but converts an exhaustive manual search into targeted review of an enriched candidate list.

### Runtime and implementation

DNA2Graph does not require training on manually annotated data and can run on a personal computer. On an Apple M1 machine with 8 GB of RAM, a 20, 000 pixels x 20, 000 pixels stitched image is processed in approximately 1-8 minutes, depending on the number of molecules present in the image. This makes the software suitable for routine use on personal computers while supporting a command-line mode for scalable execution on high-performance computing clusters.

Adaptation to new EM preparations is performed by changing interpretable parameters rather than by assembling a training set and retraining a network. This is particularly useful for rotary-shadowed DNA EM, where contrast, molecule density and background quality can vary between preparations, microscopes and imaging sessions.

## Discussion

Electron microscopy of spread DNA molecules remains a unique approach to directly visualize individual DNA replication, recombination and repair intermediates. However, meaningful conclusions require quantitative analysis of structural intermediates, which are typically rare relative to linear DNA molecules, and even enriched preparations require the visual inspection of large numbers of molecules.

DNA2Graph addresses this bottleneck by converting exhaustive visual search into targeted review of candidate structures. The software does not replace the operator’s interpretation of DNA structures; instead, it separates the large excess of linear molecules from non-linear candidates that are more likely to contain biologically informative structural intermediates.

### DNA2Graph-assisted review does not lead to missing DNA structures

Adopting DNA2Graph-assisted review does not come at the cost of missing genuine structures. In the comparative analysis of two samples with markedly different structure abundance, the rate of false negatives of DNA2Graph-assisted approach was consistently lower than that of the operator. Importantly, both operators were highly experienced in this type of analysis, and the comparisons were performed on relatively small image subsets analyzed in single, uninterrupted sessions; all conditions that favor operator performance. In typical EM experiments, structure identification would be carried out over much larger datasets and extended periods, where operator sensitivity is expected to decline with fatigue. We therefore anticipate that the sensitivity advantage of DNA2Graph-assisted review, already measurable on these curated subsets, will widen on the full-scale datasets typical of real analyses.

### False positives of DNA2Graph

In terms of software performance, false positive calls in DNA2Graph are molecules classified as non-linear that are instead judged as linear by the operator. This value ranged from ∼13 to 50% of the predicted non-linear molecules in the two samples analyzed. The main source of false positives is dirt particles adjacent to linear molecules, or areas of the grid with deposited dirt particulate. Because the operator inspects only the compact candidate list rather than the entire molecule population, discarding these false positives is rapid and does not offset the time saved. We further note that DNA2Graph defines a structure geometrically, as a physical overlap or contact between separate DNA molecules or between regions of the same molecule. Consequently, the non-linear class also includes real molecular overlaps that meet this definition but do not correspond to a biological intermediate of interest, such as chance crossings or aggregation of multiple linear molecules or imperfectly-spread DNA. These objects are correctly segmented and are not counted as false positives; rather, they are resolved by the operator during the final review of the candidate list. Distinguishing biological structure subtypes from incidental overlaps is a classification task beyond the scope of the current segmentation and is a natural application of the graph representation that DNA2Graph produces.

### Integration with ImageJ and automated downstream analysis

The consistency of the DNA2Graph output lends itself to a robust integration with ImageJ/Fiji for downstream analysis. Each molecule is assigned a consistent, human-readable identifier across all outputs generated by DNA2Graph (ROIs, length report, and spatial graph representation) allowing it to be tracked across different output types. These outputs can be directly used in ImageJ macros that automate tasks otherwise performed by hand. For example, a macro can load only the non-linear candidate ROIs and present each in turn for rapid operator classification, automating the review step across many images. In addition, because the spatial graph decomposes each candidate structure into its individual arms a macro can load these segments to measure the arm lengths of replication and recombination intermediates, such as the three arms of a fork or the four arms of a Holliday junction, enabling automated length quantification that would otherwise be tedious and error-prone by hand.

### Data handling and storage

Acquisition of large, stitched sample surfaces from EM grids for processing by DNA2Graph can lead to a rapid accumulation of imaging data. The output structure of DNA2Graph can help address this data storage burden. Once a stitch has been segmented, only the regions containing non-linear candidates (i.e. the bounding boxes of non-linear molecules) need to be retained as full-resolution EM images, while the segmentation masks and graph files provide a compact, complete record of the remaining molecules. Archiving the large original stitches while keeping these lightweight representations can reduce ongoing storage requirements substantially.

### Enabling the study of rare events

A major consequence of this shift is that DNA2Graph makes it more realistic to search for rare DNA structures that are difficult to study with conventional manual EM analysis. Rare replication intermediates, uncommon repair products and other low-frequency structures can be missed or under sampled when the number of inspected molecules is limited by operator time. By reducing the cost of screening, DNA2Graph allows larger datasets to be analyzed and should make it possible to test conditions in which the expected phenotype is a change in the frequency of a rare molecular species rather than the appearance of an abundant new structure.

### Quantitative analysis from the graph representation

The graph representation produced by DNA2Graph is not limited to structure detection but constitutes a quantitative, machine-readable description of each molecule’s contour and connectivity. This opens the way to analyses that go beyond counting structures, toward measuring geometric and physical properties of the DNA itself. Features such as contour and end-to-end length, local curvature, the presence of kinks or sharp bends, and descriptors of flexibility or persistence length can in principle be extracted directly from the graph and compared between samples or conditions. Framing single-molecule EM output as structured data rather than as images alone should make it possible to apply these quantitative and statistical methods.

### Future developments

The graph output of DNA2Graph also opens the possibility of further automation. In the present implementation, non-linear molecules are reported as candidate structures for operator review. However, the same graph representation encodes molecular connectivity, branch points, loops and contour geometry, and could therefore be used as input for future classifiers trained to recognize specific structures such as replication forks, reversed forks, termination intermediates, i-loops or circular molecules. Such classifiers could be based directly on graph-derived features, on cropped candidate images, or on hybrid approaches combining both. If trained on carefully curated datasets, this would move the analysis from automated candidate selection toward automated structure classification and would reduce operator-dependent bias in assigning molecules to specific structural classes.

The current version of DNA2Graph is therefore best viewed as a robust prescreening and measurement framework for rotary-shadowed DNA EM images. Its performance still depends on image quality, spreading conditions and parameter choice, and expert review remains necessary to reject contaminating particles, overlapping molecules or preparation-specific artifacts. Within these limits, DNA2Graph removes a major practical barrier in single-molecule DNA EM analysis and extends the range of quantitative questions that can be approached with this technique.

## Data availability

DNA2Graph is distributed via PyPI (https://pypi.org/project/dna2graph/) and its source code is available on GitHub (https://github.com/chinefed/DNA2Graph).

This paper refers to DNA2Graph v1.0.2; an archival version is available on Zenodo at https://doi.org/10.5281/zenodo.21726427.

A technical manual providing a detailed description of DNA2Graph’s internal algorithms and core logic is provided as Supplementary Data. Additional materials, including an installation guide and several video tutorials, are available on the DNA2Graph website (https://federicochinello.com/DNA2Graph/).

The subset of 20 images from the human genomic DNA dataset used to evaluate the sensitivity and false discovery rate of DNA2Graph is available on Zenodo at https://doi.org/10.5281/zenodo.21795641. DNA2Graph outputs supporting the analysis discussed in the paper, including raw CSV measurement files, segmentation masks, spatial graphs and ImageJ ROI files, will be made available through a public data-sharing repository. All other data supporting the findings of this study are included in the manuscript and its Supplementary Data.

## Supporting information

DNA2Graph Technical Manual

## Acknowledgements

We thank Martina Galli and Dana Branzei for providing the human and yeast DNA grids, respectively. We thank Cristiano Petrini and Raoul Bonnal of the IFOM Research Computing & Data Science unit for assistance with testing DNA2Graph on the IFOM high-performance computing (HPC) cluster. We thank Simonas Masiulis (Thermo Fisher Scientific) for the initial help in setting up the conditions for the tomography acquisition of rotary shadowing samples.

## Author contributions

E.Z., Y.D. and F. M. B conceptualized the project. F.C. conceived the design of DNA2Graph as a solution to the segmentation problem and developed, coded and implemented the software, including subsequent refinement in response to feedback, the user documentation and the software website. F.M.B. supervised the software development. Y.D. tested and beta-tested the software on real datasets and applied it in different experimental contexts, providing feedback throughout its development. F.B. prepared the control linear DNA sample. M.G. performed all EM spreading and image acquisition. Y.D. and F.B. performed the manual annotation as independent operators, and Y.D. carried out the quantitative analysis of the length distributions. Y.D. and F.M.B. supervised the project and acquired funding. Y.D. and F.C. wrote the manuscript, with input from all authors.

## Funding

Work in YD’s laboratory is supported by Associazione Italiana per la Ricerca sul Cancro, AIRC Grant, IG 28954 and Worldwide Cancer Research, WWCR 24-0166

## Conflict of interest

The authors declare no competing interests.

## Notes

https://federicochinello.com/DNA2Graph/

https://github.com/chinefed/DNA2Graph

### Competing Interest Statement

The authors have declared no competing interest.

