## Supplementary material for "DNA2Graph enables automated identification of non-linear DNA molecules in electron microscopy": DNA2Graph Technical Manual

Federico Chinello

Version 1.0.2 - July 31, 2026

**3 Trained Segmentation Pipeline (Beta)** **15**

**4 Supported Image Formats** **16**

**5 Length Measurement** **16**

**6 Spatial Graph Export Format** **16**

**7 Cache** **18**

### 1 Introduction

DNA2Graph is an open-source image analysis software for the automated segmentation and classification of DNA molecules in electron microscopy images obtained by rotary shadowing. It produces multiple outputs, including binary segmentation masks, ImageJ-compatible ROI annotations, quantitative length measurements, and spatial graph representations for downstream computational analysis.

This document provides a technical description of the algorithms and core logic underlying DNA2Graph. It is intended for technical users who wish to understand the software's internal functioning and configuration parameters. Throughout the document, user-adjustable parameters are denoted in `monospace font`.

### 2 Pipeline

DNA2Graph is designed as a two-stage processing framework. The pipeline, illustrated in Figure 1, consists of (i) a **Segmentation Module**, which isolates DNA structures from the background, and (ii) a graph-based **Error Correction Module**, which refines these structures to ensure continuity and biological plausibility.

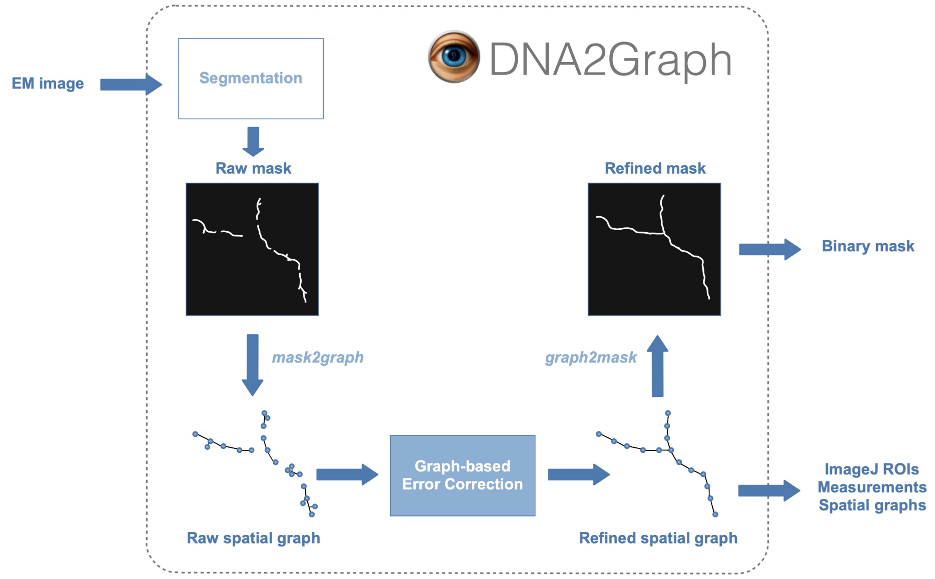

Figure 1: **DNA2Graph.** The DNA2Graph pipeline consists of a Segmentation Module that generates an initial mask, followed by a graph-based Error Correction Module that refines it.

#### 2.1 Segmentation Module

The Segmentation Module processes the input image to produce a coarse binary mask that separates DNA from the background. A high-level summary of the individual steps is provided in Table 1, while the remainder of this section presents a detailed description of each step.

Table 1: **Segmentation Module.**

| **Step** | **Purpose** |
|---|---|
| Validity mask computation | Computes a binary mask that identifies valid regions of the image, used throughout the pipeline to exclude corrupted or irrelevant areas and prevent them from influencing segmentation. |
| Preprocessing | Enhances contrast and standardizes intensity distributions to reduce inter-image variability; converts 16-bit images to 8-bit and inverts intensities so DNA molecules exhibit higher grayscale values than the background. |
| Bilateral Filter | Reduces background noise while preserving edge information, ensuring that thin DNA molecules remain well-defined for subsequent processing. |
| Unsharp Masking | Enhances edges and local contrast to better highlight DNA structures against the background. |
| Hysteresis Thresholding | Generates a binary segmentation by separating DNA molecules from the background. |
| Area-based Pruning (I) | Removes small, spurious segmented regions based on a predefined area threshold. |
| Morphological Closure | Fills small gaps and discontinuities between segmented regions to improve the structural continuity of DNA molecules. |
| Area-based Pruning (II) | Removes small, spurious segmented regions based on a predefined area threshold, which can differ from the one used in the first pruning stage. |

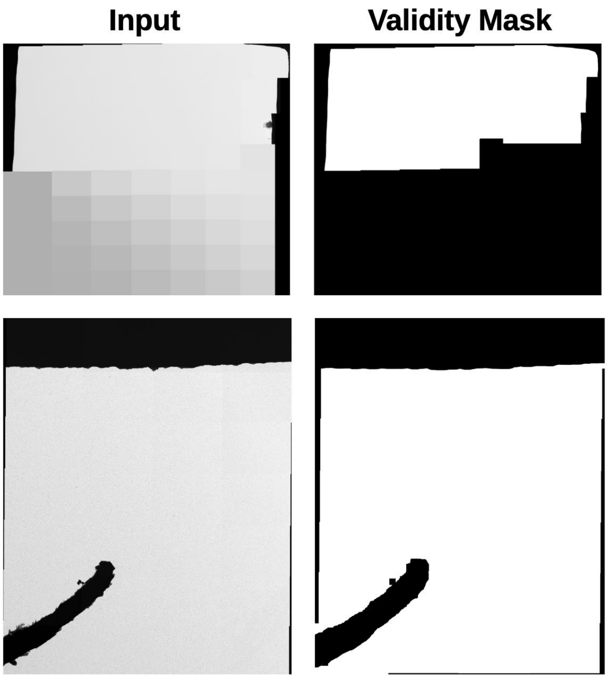

Figure 2: **Validity Mask Computation**. DNA2Graph automatically detects invalid regions in the input image and disregards them throughout the pipeline, so that they cannot influence the final output.

##### 2.1.1 Validity Mask Computation

A validity mask is computed to exclude regions outside the sample or corrupted by image acquisition errors (see Figure 2). These regions are ignored throughout the pipeline. Invalid regions are identified by low local intensity variation. A pixel is marked as invalid if the intensity range within a square neighborhood of side length `block_size` is below a `tolerance` factor times the global image intensity range. The resulting invalid regions are then expanded by `safety_margin` pixels.

##### 2.1.2 Pre-processing

Several pre-processing steps are applied:

- **Contrast enhancement** using percentile-based normalization.
- **16-bit to 8-bit conversion**, if the input image is 16-bit.
- **Histogram matching** with respect to a reference histogram obtained from a representative image.

- **Inversion**: the image complement is computed (255 - *image*).

###### 2.1.3 Bilateral Filtering

The bilateral filter is a noise-reduction technique that preserves edges by combining spatial proximity and intensity similarity. This allows smoothing of homogeneous regions while maintaining sharp boundaries.

- `bilateral_diameter`: Diameter of the pixel neighborhood used during filtering.
- `bilateral_sigma_color`: Standard deviation in the intensity domain; controls how dissimilar colors are mixed.
- `bilateral_sigma_space`: Standard deviation in the spatial domain; controls the influence of distant pixels.

###### 2.1.4 Unsharp Masking

Unsharp masking enhances image sharpness and improves local contrast by amplifying high-frequency components.

- `high_boost_radius`: Standard deviation of the Gaussian blur applied before detail extraction.
- `high_boost_amount`: Scaling factor controlling the strength of the sharpening effect.

###### 2.1.5 Hysteresis Thresholding

This step binarizes the image using a dual-threshold approach. The lower threshold $L$ and upper threshold $H$ are computed from percentiles of valid pixel intensities.

- $L$: `hysteresis_low_pct` percentile of valid intensities.
- $H$: `hysteresis_high_pct` percentile of valid intensities.

Pixels with intensity above $H$ are marked as strong, those below $L$ are discarded, and intermediate pixels are retained only if connected (directly or indirectly) to strong pixels.

###### 2.1.6 Area-based Pruning (I)

Connected components with area smaller than a predefined threshold are removed.

- `min_area_1`: Minimum area required for a component to be retained.

###### 2.1.7 Morphological Closure

Morphological closing is applied to fill small holes and connect nearby regions.

- `closure_kernel_size`: Size of the structuring element used for the closing operation.

###### 2.1.8 Area-based Pruning (II)

A second pruning step removes remaining small components after morphological processing.

- `min_area_2`: Minimum area required for a component to be retained.

#### 2.2 Error Correction Module

The binary mask produced by the Segmentation Module is used to extract a spatial graph representation of the DNA molecules. This spatial graph is then refined by an Error Correction Module consisting of a cascade of custom graph-processing algorithms. These algorithms are summarized at a high level in Table 2 and described in detail in the remainder of this section. Finally, the cleaned graph representation is converted back into image form.

Table 2: **Error Correction Module.**

| **Step** | **Purpose** |
|---|---|
| Mask-to-Graph Conversion | Converts each segmented DNA molecule in the mask into a spatial graph representation, where nodes correspond to pixels along the molecule with their $(x, y)$ coordinates, and edges encode connectivity. |
| Repair Connectivity (I) | Operates on spatial graph representations to address fragmentation caused by segmentation errors or signal dropout in thin DNA structures, by adding edges between relevant node pairs to restore connectivity. |
| Prune Short Cycles | Operates on spatial graph representations to remove spurious short cycles, using a predefined cycle-length threshold. |
| Prune Short Branches | Operates on spatial graph representations to remove spurious short branches, using a predefined branch-length threshold. |
| Prune by Thickness | Cleans the spatial graph representations by removing nodes associated with abnormally thick regions, leveraging the prior that DNA structures are thin and thicker regions are typically spurious. |
| Prune Small Components | Discards small spatial graph representations that are likely spurious. |
| Repair Connectivity (II) | Further refines spatial graph representations by performing a second connectivity repair step, using different parameters from the first repair stage. |
| Graph-to-Mask Conversion | Converts the refined spatial graph representations back into mask form. |

##### 2.2.1 Mask-to-Graph Conversion

A spatial graph representation is extracted from the binary mask generated by the Segmentation Module. The mask is first skeletonized using the Zhang-Suen thinning algorithm. Skeletonization yields a one-pixel-wide representation that preserves the shape and connectivity of each DNA molecule. The skeletonized mask is then converted into a spatial graph $G = (V, C, E)$, where $V$ is the set of nodes, $C$ denotes their spatial coordinates, and $E$ is the set of edges connecting node pairs $(u, v) \in V$.

Graph construction proceeds by mapping each skeleton pixel to a node, with its $(x, y)$ image coordinates[1] stored as spatial attributes (Figure 3). Edges are introduced between nodes whose corresponding pixels are immediate neighbors in the image grid, following the standard notion of 8-connectivity. Each molecule is represented as a connected component in the spatial graph.

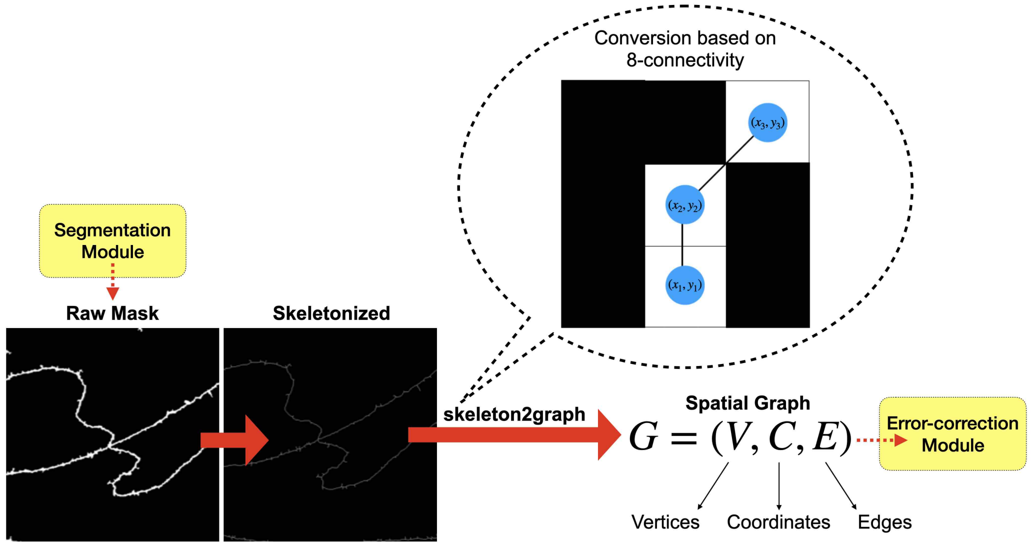

Figure 3: **Skeleton-to-graph Conversion.** To convert a skeleton into a spatial graph, each pixel that lies on the skeleton is mapped to a corresponding node in the graph, with its $(x, y)$ coordinates stored as spatial attributes. An undirected edge is added between two nodes if their corresponding pixels in the skeleton are 8-connected, i.e., adjacent in the horizontal, vertical, or diagonal direction.

[1] DNA2Graph internally represents coordinates as $(y, x)$. Throughout this manual, for simplicity, we adopt the $(x, y)$ convention.

##### 2.2.2 Repair Connectivity (I)

The algorithm restores connectivity in the spatial graph representation by adding edges between nodes that are close in Euclidean space but far apart in graph distance. It operates on endpoint nodes (i.e., nodes of degree 1) as sources, since these are natural candidates for broken connections.

For each endpoint, a spatial neighborhood is constructed as the set of all nodes within a given Euclidean radius, forming an initial candidate set. This set is then filtered to retain only nodes whose shortest-path distance from the endpoint exceeds a specified threshold. If the resulting set is non-empty, the endpoint is connected to the candidate that is closest in Euclidean space.

Optionally, the candidate set can be restricted to endpoint nodes only, so that connections are introduced exclusively between endpoints.

**Examples:**

- *Disconnected components.* An endpoint $e$ and a node $v$ are spatially close (their Euclidean distance is within $d_e$) and belong to distinct connected components (their shortest-path distance is $\infty$). An edge $(e, v)$ is added, merging the two components.

- *Topologically close nodes.* An endpoint $e$ and a node $v$ are spatially close (their Euclidean distance is within $d_e$) and topologically close (their shortest-path distance is within $d_g$). No edge is added.

##### 2.2.3 Prune Short Cycles

The algorithm simplifies the spatial graph representation by iteratively contracting short cycles. At each iteration, it first computes a cycle basis and selects the cycles whose length is below a prescribed threshold. These short cycles are then processed one at a time, ensuring that overlapping cycles are not modified within the same iteration. Each cycle is replaced by a single node, whose spatial coordinates are defined as the centroid of the coordinates of its neighboring nodes. If a cycle has no neighboring nodes, it is simply removed. If replacing a cycle would introduce a node with excessively high degree, the entire component is considered unreliable and is discarded.

The procedure is repeated until no further contractions are possible or until a maximum number of iterations, specified as a parameter of the algorithm, is reached. The algorithm operates independently on each connected component, since different components may require a different number of iterations.

**Algorithm 1** Repair Connectivity

**Require:** Spatial graph $G = (V, E)$
**Require:** Euclidean distance threshold $d_e$ (`max_euclidean_distance`)
**Require:** Graph-distance threshold $d_g$ (`min_graph_distance`)
**Require:** Endpoint restriction flag (`restrict_to_endpoints`)

$S \leftarrow \{ e \in V \mid \deg(e) = 1 \}$ ▷ Sources
**if** endpoint restriction is enabled **then**
  $C \leftarrow S$
**else**
  $C \leftarrow V$
**end if**
**for all** $e \in S$ **do**
  $N_e \leftarrow \{ v \in C \mid \|e - v\| \leq d_e \}$ ▷ Euclidean neighborhood
  $T_e \leftarrow \{ v \in N_e \mid \mathrm{dist}_G(e, v) \geq d_g \}$ ▷ Graph-distance filtering
  **if** $T_e \neq \emptyset$ **then**
    $v^* \leftarrow \arg\min_{v \in T_e} \|e - v\|$
    $E \leftarrow E \cup \{(e, v^*)\}$
  **end if**
**end for**
**return** $G$

**Algorithm 2** Prune Short Cycles

**Require:** Spatial graph $G = (V, E)$
**Require:** Minimum cycle length $\ell_{\min}$ (`min_cycle_length`)
**Require:** Maximum number of iterations $k_{\max}$
**Require:** Wiring cap $w_{\max}$

```
Decompose $G$ into connected components $\mathcal{C} = \{G_1, \dots, G_m\}$
for all $G_i \in \mathcal{C}$ do
    for $k = 1$ to $k_{\max}$ do
        $B \leftarrow$ a cycle basis of $G_i$
        $S \leftarrow \{\, c \in B \mid |c| < \ell_{\min} \,\}$                ▷ Short cycles
        if $S = \emptyset$ then
            break
        end if
        $R \leftarrow \emptyset$                                          ▷ Nodes removed
        for all $c \in S$ do
            if $c \cap R \neq \emptyset$ then
                continue                                      ▷ Skip overlapping cycles
            end if
            $N_c \leftarrow \{\, v \in V(G_i) \setminus c \mid \exists\, u \in c$ with $(u, v) \in E(G_i) \,\}$
            Remove all nodes in $c$ from $G_i$
            $R \leftarrow R \cup c$
            if $N_c = \emptyset$ then
                continue                                      ▷ Discard isolated cycle
            end if
            if $|N_c| > w_{\max}$ then
                Clear $G_i$                                   ▷ Discard corrupted component
                break
            end if
            $v^* \leftarrow \mathrm{centroid}(N_c)$
            Add $v^*$ to $G_i$
            Add edges $\{\, (v^*, v) \mid v \in N_c \,\}$ to $G_i$
        end for
    end for
end for
$G \leftarrow$ recomposition of all components in $\mathcal{C}$
return $G$
```

##### 2.2.4 Prune Short Branches

The algorithm simplifies the spatial graph representation by iteratively pruning short branches. A branch is defined as a maximal path starting from an endpoint and traversing only nodes of degree 2, until a node of degree different from 2 is reached. This terminal node is referred to as the *joint* of the branch, i.e., the node at which the branch attaches to the rest of the graph.

The algorithm proceeds iteratively: at each iteration, it identifies all branches whose length is below a prescribed threshold. These short branches are then processed in increasing order of length, so that the shortest and most likely spurious branches are removed first. The algorithm prevents the removal of multiple branches that share the same joint within a single iteration. This restriction is essential because a branch that is classified as short at the beginning of an iteration may no longer satisfy this condition after a neighboring branch has been removed.

For instance, consider a Y-shaped structure in which two short branches meet at a joint, while a third segment (the "trunk" of the Y) connects the joint to the rest of the graph. If one of the two short branches is removed, the joint loses one incident edge and its degree drops from 3 to 2, so that it is no longer a junction. As a result, the remaining short branch no longer terminates at that node, but instead continues through it and forms a longer path together with the trunk; consequently, it may no longer be classified as short. For this reason, once a branch incident to a given joint is removed, all other branches attached to that joint are deferred to the next iteration, where they are re-extracted and re-evaluated on the updated graph.

The procedure continues until there are no remaining short branches or until a maximum number of iterations, specified as a parameter of the algorithm, is reached. The algorithm operates independently on each connected component, since different components may require a different number of iterations.

**Algorithm 3** Prune Short Branches

**Require:** Spatial graph $G = (V, E)$
**Require:** Minimum branch length $\ell_{\min}$ (`min_branch_length`)
**Require:** Maximum number of iterations $k_{\max}$

```
Decompose $G$ into connected components $\mathcal{C} = \{G_1, \ldots, G_m\}$
for all $G_i \in \mathcal{C}$ do
    for $k = 1$ to $k_{\max}$ do
        $S \leftarrow \{ v \in V(G_i) \mid \deg(v) = 1 \}$        ▷ Endpoints
        $B \leftarrow \emptyset$        ▷ Branches marked for removal
        for all $e \in S$ do
            Extract the branch $b_e$ starting at $e$
            if $|b_e| < \ell_{\min}$ then
                $B \leftarrow B \cup \{b_e\}$
            end if
        end for
        if $B = \emptyset$ then
            break
        end if
        $J \leftarrow \emptyset$        ▷ Processed joints
        for all $b \in B$ in increasing order of length do
            $v_b \leftarrow$ joint of $b$
            if $v_b \notin J$ then
                Remove all nodes of $b$ except $v_b$
                $J \leftarrow J \cup \{v_b\}$
            end if
        end for
    end for
end for
Recombine all components into $G$
return $G$
```

##### 2.2.5 Prune by Thickness

DNA molecules are thin filaments and thick regions in the mask produced by the Segmentation Module are typically associated with noise or artifacts. Each node/pixel in the spatial graph representation carries an attribute which captures the approximate thickness radius of the binary mask at that pixel. The Prune by Thickness algorithm simplifies the spatial graph by removing nodes that exhibit excessive thickness.

Nodes whose thickness radius exceeds a prescribed `max_thickness` threshold are considered candidates for removal. However, such nodes are only removed if they are not structurally important, as their removal could otherwise disconnect relevant parts of the graph.

A node is defined as structurally important if it plays a significant role in the connectivity of the graph. This is the case if either:

- the node belongs to a cycle; or
- the node has at least two structurally important incident edges.

More precisely, an incident edge is considered structurally important if, when followed away from the node, it either:

- reaches nodes at a distance greater than a prescribed depth cutoff (`structurality_depth_cutoff`), indicating that it connects to a sufficiently extended structure; or
- leads to a cycle anywhere in the graph, indicating connection to a loop-like structure.

Requiring at least two such edges ensures that the node is not merely attached to a single significant structure, but instead acts as a connector between multiple meaningful regions of the graph. Nodes that do not satisfy these conditions are considered non-structural and can be safely removed if their thickness radius exceeds the prescribed threshold.

##### 2.2.6 Prune Small Components

The spatial graph representation is pruned by removing connected components whose largest bounding box side is smaller than the user-defined threshold `minimum_bbox_size`, thereby discarding small and likely spurious structures. Here, the bounding box is defined as the smallest axis-aligned rectangle enclosing all nodes of a connected component.

##### 2.2.7 Repair Connectivity (II)

The spatial graph representation is further refined by a second pass of the Repair Connectivity algorithm, using a different set of parameters from the first pass.

##### 2.2.8 Graph-to-Mask Conversion

Once error correction is complete, the cleaned spatial graph representation is rendered back onto the pixel grid, yielding a binary segmentation mask that retains the improvements from graph-based repair. The conversion first rasterizes a line between the endpoints of every graph edge, producing intermediate pixels for long edges added during connectivity repair. It then applies a $5 \times 5$ dilation.

### 3 Trained Segmentation Pipeline (Beta)

DNA2Graph includes an optional analysis mode that replaces its standard Segmentation Module (Section 2.1) with a custom CNN U-Net model (`Use Trained Segmentation Pipeline`). The U-Net was trained from scratch using segmentation masks generated automatically by DNA2Graph itself, without manual annotation.

Specifically, standard, non-learning-based DNA2Graph was first applied to a training dataset to produce pseudo-labels, which were then used as target masks for training the U-Net. In this teacher–student framework, DNA2Graph acts as the "teacher," while the U-Net serves as the "student". Although the model is trained using labelled targets, these labels are generated automatically; therefore, the training process does not require manually annotated data.

When the U-Net is enabled, its output is subsequently processed using a simplified graph-based Error Correction Module.

The trained segmentation pipeline is currently provided as a beta feature, and we recommend that most users employ the standard segmentation pipeline for their analyses.

### 4 Supported Image Formats

DNA2Graph supports 8-bit and 16-bit grayscale images in TIFF format. The software was thoroughly tested on images acquired at pixel sizes of approximately 1.0–1.7 nm/pixel. Large deviations from this range may require parameter re-tuning for optimal performance.

### 5 Length Measurement

DNA2Graph can export a report containing length measurements for individual molecules. Lengths are expressed in pixels and can be converted to the desired unit (e.g., nanometers) by multiplying each length by the physical pixel size. For example: length (nm) = length (pixels) × pixel size (nm/pixel).

### 6 Spatial Graph Export Format

The cleaned spatial graph representation generated by the Error Correction Module is exported in a custom HDF5 format, where each connected component, corresponding to a molecule, is stored as a separate group.

A connected component / molecule is represented as a set of *linear walks*, defined as maximal non-branching paths. Each walk is an ordered sequence of nodes in which all intermediate nodes have degree two (node $n$ is connected to node $n + 1$ by an edge). Walks are encoded through two datasets:

- The `coords` dataset is a 2D array of shape $(N, 2)$ containing the concatenated $(y, x)$ coordinates of all walks.
- The `offsets` dataset is a 1D array of length $K + 1$ that partitions `coords` into $K$ individual walks, such that the $i$-th walk is given by `coords[offsets[i]:offsets[i+1]]`.

The component / molecule label (non-linear, linear, or boundary) is stored as a group attribute.

A molecule is reconstructed as a set of linear walks, i.e., coordinate sequences, by slicing `coords` according to `offsets`:

```
import h5py
import numpy as np

def load_molecule(h5_grp):
        coords = h5_grp['coords'][:]
        offsets = h5_grp['offsets'][:]

        seqs = []
        for i in range(len(offsets) - 1):
                start, end = offsets[i], offsets[i+1]
                seq = coords[start:end]
                seqs.append(seq)

        return seqs
```

Adjacency between walks is recovered by identifying sequences that share the same terminal coordinates:

```
from collections import defaultdict
import numpy as np

def build_adjacency(seqs):
        n_seqs = len(seqs)
        adjacency = np.zeros((n_seqs, n_seqs))

        terminal_map = defaultdict(list)
        for i, seq in enumerate(seqs):
                first = tuple(int(x) for x in seq[0].tolist())
                last = tuple(int(x) for x in seq[-1].tolist())
                terminal_map[first] += [i]
                terminal_map[last] += [i]

        for indices in terminal_map.values():
                for i in range(len(indices)):
                        for j in range(i + 1, len(indices)):
                                adjacency[indices[i], indices[j]] = 1
                                adjacency[indices[j], indices[i]] = 1

        return adjacency
```

The walks can be easily rendered into a binary mask. This is achieved by initializing an empty image canvas and assigning foreground values to the pixel locations corresponding to the coordinates of the graph nodes:

```
import cv2
import numpy as np

def render_walks(seqs):
        all_points = np.concatenate(seqs)
        min_y, min_x = np.min(all_points, axis=0)
        max_y, max_x = np.max(all_points, axis=0)
        width = max_x - min_x + 1
        height = max_y - min_y + 1

        canvas = np.zeros((height, width), dtype=np.uint8)
        y = all_points[:, 0] - min_y
        x = all_points[:, 1] - min_x
        canvas[y, x] = 255

        dilation_kernel = np.ones((5, 5), np.uint8)
        canvas = cv2.dilate(canvas, dilation_kernel, iterations=1)

        return canvas
```

Note that this simple function is aimed for quick inspection and, differently from the rendering approach used internally by DNA2Graph (Section 2.2.8), it does not add intermediate pixels along the long edges introduced by connectivity repair. As a result, the rendered molecule may display gaps even if the underlying graph is connected.

### 7 Cache

DNA2Graph caches the spatial graph representation generated immediately before the Prune Small Components step of the Error Correction Module. The graph is stored in Python pickle format as `.cached_graph.pkl`, within a hidden `.cache` directory in the image output folder.

This cache allows users to modify the parameters of the Prune Small Components and Repair Connectivity (II) steps without rerunning all preceding stages of the analysis. To perform a clean analysis without reusing previously cached data, enable the `Clean Cache` option.
